# Ablation of microglial estrogen receptor alpha predisposes to diet-induced obesity in male mice

**DOI:** 10.1101/2025.11.03.685977

**Authors:** Inmaculada Velasco, Jeremy M. Frey, Vladislav Baglaev, Tristan Jafari, Thomas Huang, Sophia Schwie, Olivia D. Santiago, Rachael D. Fasnacht, Joshua P. Thaler, Mauricio D. Dorfman

## Abstract

Estrogen receptor alpha (ERα) signaling has metabolic and anti-inflammatory properties in addition to its impact on reproductive function. In male but not female mice, inflammatory activation of microglia, the resident macrophages of the brain, has been implicated in the pathogenesis of diet-induced obesity (DIO), raising the possibility that differences in microglial estrogen signaling may account for the sexual dimorphism. In this study, we assessed metabolic and CNS histopathological properties in a mouse model with inducible microglia-specific ablation of ERα (MG-ERαKO). Male MG-ERαKO mice developed increased weight gain and insulin resistance relative to controls during high-fat diet (HFD) feeding. Indirect calorimetry analysis revealed that reduced energy expenditure was the main driver of the obese phenotype. In contrast, female MG-ERαKO mice fed HFD developed mild insulin resistance with no change in body weight gain compared to controls. Immunohistochemical analyses of the microglial activation marker IBA1 in the mediobasal hypothalamus (MBH) revealed that female MG-ERαKO mice had an increased number of microglia without showing morphological signs of activation. In contrast, MBH microglial number was unchanged in MG-ERαKO male mice, but the cells adopted more activated morphological profiles. Finally, HFD-fed MG-ERαKO male mice had increased POMC neuron-microglia interactions but fewer overall hypothalamic POMC neurons, suggesting microglia may disrupt POMC neuron integrity to promote DIO. Together, these findings indicate that sex-specific actions of estrogen in microglia limit the metabolic complications of HFD feeding.

## 1. Introduction

Estrogen is the archetypal female sex steroid hormone with a critical role in reproductive function. However, sex steroids also have pleiotropic actions throughout different body systems, including modulating energy metabolism and immune function [1,2]. Estrogen deficiency associated with menopause or induced surgically (ovariectomy (OVX)) or pharmacologically (inhibition of aromatase, the key enzyme that generates E2 from testosterone) leads to increased adiposity, reduced energy expenditure, and increased tissue inflammation [3–5]. Likewise, in men, the aging-related decline in testosterone leads to lower circulating levels of estrogen, again associated with increased adiposity [1,6]. Mechanistically, estradiol (E2; the main circulating estrogen) acts predominantly through estrogen receptor (ER) alpha (ERα) [7–9]. Indeed, mice with global ablation of ERα or lacking ERα in the CNS alone are obese and insulin resistant due to reduced energy expenditure and increased food intake [10–13]. However, deletion of ERα from specific populations of hypothalamic neurons, including pro-opiomelanocortin (POMC) neurons, affects energy balance and diet-induce obesity (DIO) sensitivity only in females, raising the possibility that other CNS cell types mediate estrogen actions in males. Regarding the role of estrogen in suppressing inflammatory signaling, evidence from neurological diseases such as multiple sclerosis, Alzheimer’s disease, and Parkinson’s disease shows that estrogens alleviate disease progression by reducing neuroinflammation and glial cell activation [14–16], supporting a role of ER signaling in non-neuronal cells during DIO.

DIO is linked to dysfunction of hypothalamic POMC neurons [17–20], which regulate energy balance by controlling food intake and energy expenditure [17]. Increasing evidence suggests that hypothalamic inflammation and glial activation contribute to this neuronal impairment [20–26]. As the innate immune cells of the CNS, microglia respond to combat infection and repair injury but are also implicated in neurologic diseases. During these events, microglia undergo a characteristic morphological transformation from a homeostatic state with long processes extending from the soma to an “activated” profile characterized by a more rounded, ameboid shape with retracted and thickened ramifications [27]. Aside from established roles in maintenance of CNS immune homeostasis and postulated contributions to neurodegenerative disease progression, microglia also contribute to obesity pathogenesis in rodents and humans [22,23,28–31]. Histopathologic analyses performed postmortem and in vivo neuroimaging support the presence of inflammation and gliosis in the mediobasal hypothalamus (MBH) of children and adults with obesity [32]. In animal models of DIO, hypothalamic microglial activation is observed prior to substantial weight gain within days of high-fat diet (HFD) feeding onset [22]. Pharmacologic and genetic interventions targeting microglia show that reduction of microglial activation and inflammation induced by HFD feeding protects from DIO [20,24,25,30,33,34], and prevents POMC neuron dysfunction [20,24–26,35]. Importantly, female mice exposed to HFD develop minimal hypothalamic microgliosis and inflammation compared to males [31,36], a sex difference that disappears in estrogen-deficient OVX animals [37]. Furthermore, E2 administration to OVX mice prevents the acute hypothalamic microgliosis induced by short-term HFD feeding [38], suggesting the possibility that estrogen signaling in microglia protects from DIO and hypothalamic gliosis.

While several studies using ER KO mice, microglial cell lines, and primary microglial cultures have demonstrated direct anti-inflammatory effects of E2 on microglia [9,39–41], *in vivo* studies using microglia-specific models to determine the functional relevance of these observations are lacking. In this study, we report the role of microglia ERα signaling in protecting from HFD-induced obesity and hypothalamic microgliosis by characterizing a novel inducible microglial-specific ERα KO mouse.

## 2. Material and Methods

### 2.1. Animals and diet

Microglial-specific estrogen receptor α knockout mice (MG-ERαKO) were generated by the following breeding strategy. Heterozygous Cx3cr1^CreER^ mice (JAX strain #020940) were bred with homozygous ERα^fl/fl^ mice (kindly donated by Dr. James R. Sowers, with permission from Pierre Chambon) [42]. Experimental mice were Cx3cr1^CreER/+^::ERα^fl/fl^ (MG-ERαKO) and Cx3cr1^+/+^::ERα^fl/fl^ littermate controls (Ctl) (Figure 1A). In this model, ERα gene deletion occurs via recombination of loxP sites surrounding exon 3 of *Esr1* (ERα gene) after adult mice (6-7 weeks of age) are exposed to tamoxifen (4 consecutive days of 2 mg intraperitoneal (i.p.) injections in purified corn oil (20 mg/ml, T5648; Sigma-Aldrich)). Littermate control mice received the same tamoxifen regimen to ensure equivalent exposure. While a 4-week period is necessary to allow for turnover and replacement of CX3CR1-expressing peripheral myeloid cells by newly born non-recombined monocytes derived from the bone marrow [32], an additional 4-week period post-tamoxifen (8 weeks total) was added before any experiments and diet treatments were performed to ensure the washout of tamoxifen. Microglia are long-lived and replenished through clonal proliferation, so they retain the recombined allele indefinitely [43,44]. Male and female Ctl and MG-ERαKO mice were kept group-housed with ad libitum access to food and water, except during indirect calorimetry experiments. All mice were switched to a HFD (60% kcal from fat, D12492i; Research Diets) for 15 weeks until study termination. Cages were placed in temperature-controlled rooms with a 14:10-h light-dark cycle under specific pathogen-free conditions. All procedures were performed in accordance with *NIH Guide for the Care and Use of Animals* and were approved by the Institutional Animal Care and Use Committee at the University of Washington.

**Figure 1.**
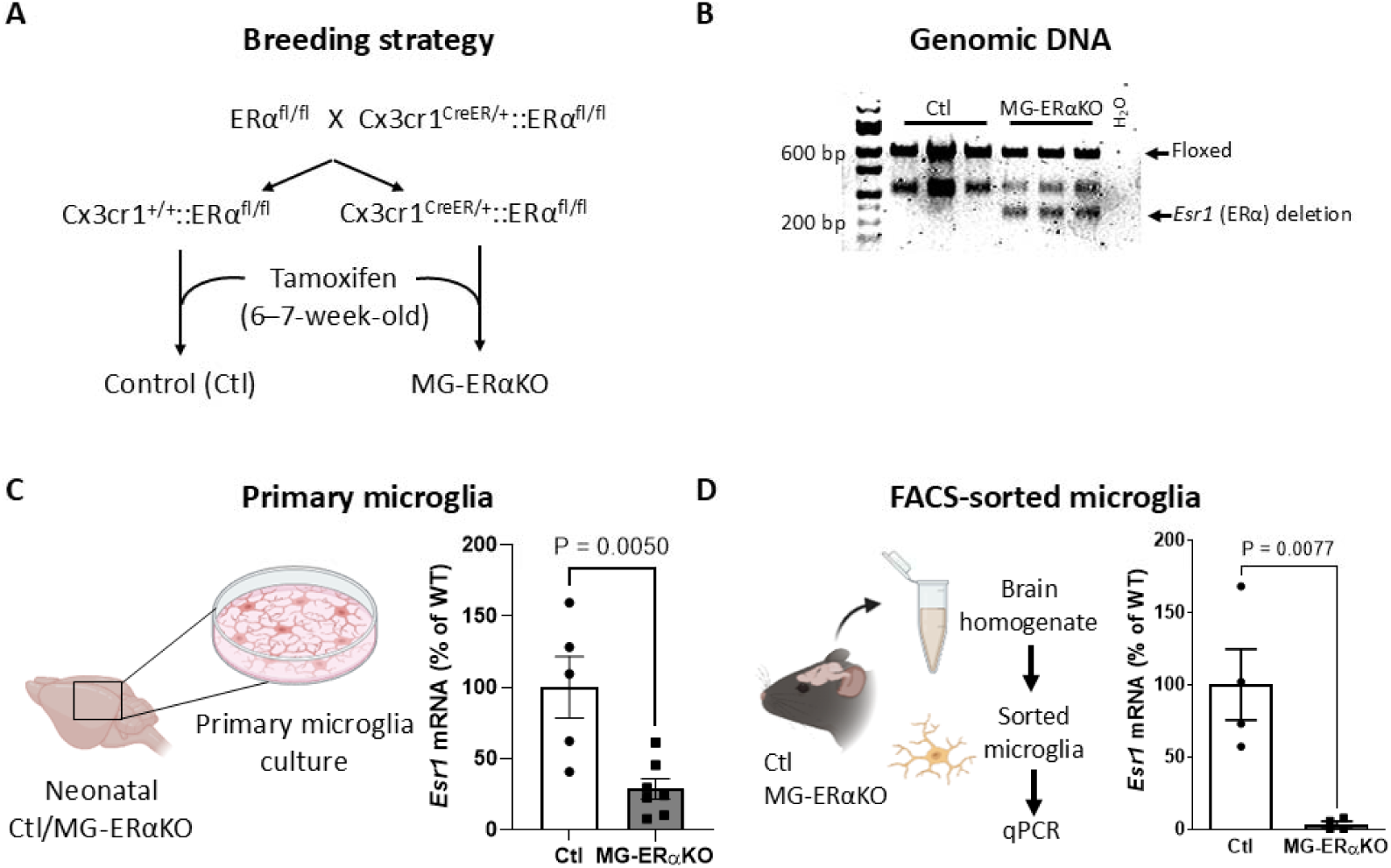
**Breeding strategy and validation of MG-ER**_α_**KO mouse.** (A) Schematic of the breeding strategy used to generate Cx3cr1-specific estrogen receptor α (ERα; encoded by the *Esr1* gene) deletion by crossing ERα^fl/fl^ mice with tamoxifen-inducible Cx3cr1-Cre^ER^ mice. (B) Recombinant PCR band indicating *Esr1* deletion, detected only in hypothalamic genomic DNA from MG-ERαKO but not control (Ctl) mice. H_2_O = no template control, (C-D) *Esr1* mRNA expression in primary microglia cultures (C) and brain-isolated microglia (D) from MG-ERαKO mice compared with Ctl littermates. Data are presented as mean ± SEM. Data was analysed by Student’s *t*-test.

### 2.2. Genotyping

Cx3cr1^CreER^ and ERα^fl/fl^ mice were genotyped by PCR using ear genomic DNA with the following set of primers: for CreER or wild-type allele - *Forward*: AAG ACT CAC GTG GAC CTG CT, *Mutant Reverse*: CGG TTA TTC AAC TTG CAC CA, *Wild Type Reverse*: AGG ATG TTG ACT TCC GAG TTG; for ERα flox or wild-type allele - *Forward*: TTG CCC GAT AAC AAT AAC AT, *Mutant Forward*: GTG TCA GAA AGA GAC AAT, *Reverse*: GGC ATT ACC ACT TCT CCT GGG AGT CT. The PCR reaction was performed with EconoTaq PLUS (30033-1; Biosearch Technologies). Amplicon sizes were: 300 bp for the CreER knock-in and 695 bp for the wildtype control (data not shown); 607 bp for the ERα floxed allele and ∼250 bp for the deletion of exon 3 (Figure 1B).

### 2.3. Experimental design

Ctl and MG-ERαKO mice of both sexes were fed with HFD for 15 weeks. Body weight (BW) and food intake (FI) were monitored weekly from week 1 to 11 during HFD feeding. Food intake was approximated by dividing the total food consumed by the number of mice in each cage (4-5). Between weeks 11 and 15, mice were subjected to glucose and insulin tolerance tests (GTT and ITT), a glucose-stimulated insulin secretion (GSIS) test, and body composition (only males). A subset of male mice (8 per group) was placed on metabolic cages to determine food intake, energy expenditure, and locomotor activity.

### 2.4. Glucose and insulin tolerance test

GTTs and ITTs were conducted at week 10 and 11 of HFD, respectively, in 4-h-fasted mice by i.p. injections of 30% D-glucose (2 g/kg; Hospira, NDC 00409-6648-02) or insulin (0.75 U/kg; Lilly, NDC 0002-8215-17). Blood glucose levels were determined at 0 (baseline), 15, 30, 60, 90 and 120 m post injection in tail capillary blood applied to a handheld glucose meter (Freestyle Lite; Abbott). Total area under the curve (AUC) was calculated by the trapezoid rule.

### 2.5. Glucose-stimulated insulin release

GSIS was performed at 12 weeks of HFD feeding in 5-h-fasted mice by i.p. injection of glucose (30% D-glucose; 2 g/kg). Tail blood samples were collected before (time 0) and 15 min post-glucose injection, using heparinized coated capillary tubes (VWR 15401-560). Blood samples were centrifuged (10 min, 2000 g, 4°C), and plasmas were collected and stored at −80 °C. Glucose levels were determined at time 0 and 15 min using a handheld glucose meter (Freestyle Lite; Abbott). Insulin was measured by ELISA (Crystal Chem 90080). Homeostatic Model Assessment for Insulin Resistance index (HOMA-IR) was calculated according to the formula: time 0 glucose (mg/dL) x time 0 insulin (ng/mL)/22.5 [45].

### 2.6. Body composition

Determination of body fat mass and lean mass was performed using quantitative magnetic resonance spectroscopy (EchoMRI, Houston, TX) in conscious mice performed by the Energy Balance Core of the NIH-funded Nutrition Obesity Research Center (NORC) at the University of Washington [46].

### 2.7. Indirect calorimetry

A subset of 8 mice per group (Ctl and MG-ErαKO) were single-housed and acclimated to metabolic cages before measurements of food intake, energy expenditure using a computer-controlled indirect calorimetry system (Promethion®, Sable Systems, NV) in the NORC Energy Balance Core at the University of Washington. O_2_ consumption and CO_2_ production were measured for each animal for 1 min at 10-min intervals as previously described [47,48]. Respiratory quotient was calculated as the ratio of CO_2_ production to O_2_ consumption, and energy expenditure was calculated using the Weir equation followed by ANCOVA analysis [49]. Ambulatory activity was measured continuously with consecutive adjacent infrared beam breaks in the x-, y- and z-axes scored as an activity count that was recorded every 10 min [47,48]. Data acquisition and instrument control were coordinated by MetaScreen v.1.6.2 and raw data was processed using ExpeData v.1.4.3 (Sable Systems) using an analysis script documenting all aspects of data transformation. Light and dark cycle energy expenditure values are reported based on averaging 72 data points per 12 h cycle on 3 consecutive days, and these in turn were averaged to obtain total 24 h energy expenditure.

### 2.8. Microglia primary cultures

Microglia primary cultures were prepared as described previously [30]. Briefly, cerebral cortices were harvested from Ctl and MG-ERαKO mice at postnatal day 1-2 (P1-2), minced and triturated in Dulbecco’s Modified Eagle’s Medium (DMEM), 4.5 g/L glucose, 10% F12 Supplement, 10% horse serum, l-glutamine, 10 mM HEPES, and 1% penicillin/streptomycin. Mixed cortical cells were seeded onto a poly-d-lysine coated T75 flask and incubated at 37 °C and 5% CO_2_/air. After 7-10 days in culture media, microglia were removed by shaking at 200 rpm for 1 h, washed with DPBS, and plated onto 24-well plates. To induce recombination, cells were treated with 5 μM of 4-hydroxytamoxifen (4-OHT; Sigma) for 48 hours. Cells were then washed twice with DPBS and lysed for RNA extraction and subsequent analyses.

### 2.9. Microglial isolation from adult mice

Microglia were isolated from whole brains of Ctl and MG-ERαKO mice to determine the efficiency of recombination [50]. Briefly, mice were transcardially perfused with PBS and brains were dissected and homogenized using Potter-Elvehjem tissue homogenizers (Wheaton), and then samples were centrifuged for 10 min at 4°C at 900 g. Cells were resuspended in 33% Percoll (Cytiva) and a PBS layer was carefully added to each tube and then samples were centrifuged for 15 min at room temp at 400 g with minimal acceleration and braking. Supernatant and myelin layers were removed, and cell pellets were resuspended and washed in fluorescence activated cell sorting (FACS) buffer (2% fetal bovine serum in PBS) for 5 min. Samples were pelleted, resuspended, passed through a 70-100 μM filter, and centrifuged for 10 min at 4°C at 400 g. Pellets were resuspended by vortexing, and simultaneously stained for CD11b and Fc-blocked on ice for 20 min (1:1:48, CD11b:Fc blocker:FACS buffer; Bio-Rad). Samples were washed 2x more with FACS buffer to remove excess antibody. Cells were stained with DAPI (1:50,000; Thermo Fisher) and immediately isolated by FACS for the live (DAPIlo) myeloid (CD11b+) population. Sorted cells were collected in RNA lysis buffer (Qiagen) with 1% β-mercaptoethanol (Sigma) and subsequently processed for qRT-PCR.

### 2.10. Real-Time PCR

Total RNA was extracted using RNeasy micro kit according to the manufacturer’s instructions (Qiagen) and reverse-transcribed with Multiscribe Reverse Transcriptase (Applied Biosystems). Levels of mRNA for *Esr1* (ERα gene), inflammatory and metabolic genes, and *Rn18s* (internal control) were measured by semi-quantitative real-time PCR (40 cycles of 95°C for 15 s and 60°C for 1 min) on an ABI Prism 7900HT (Applied Biosystems) with SYBR Green PCR Supermix (Bio-Rad). Primer sequences are listed in Table 1. The relative gene expression was calculated by the 2^-ΔΔC^_T_ method, where C_T_ is threshold cycle [51].

**Table 1.**
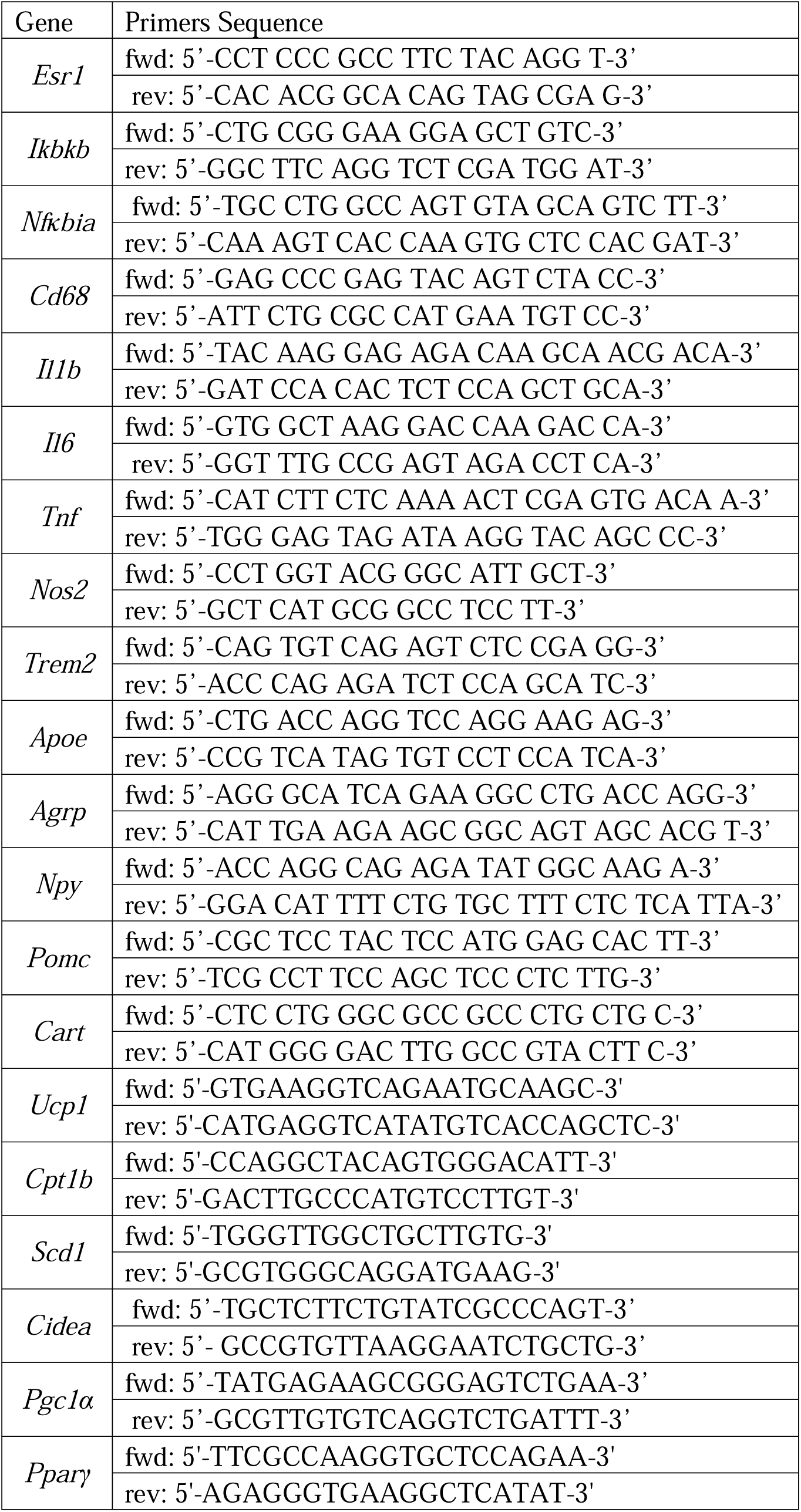
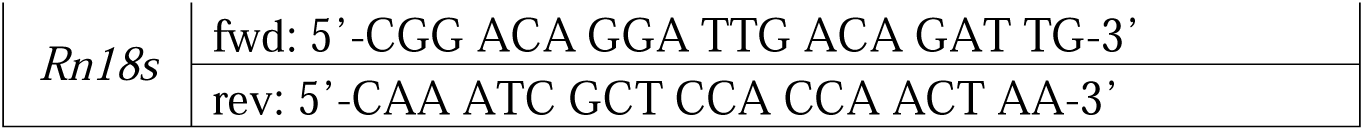
Real-time qPCR primers.

### 2.11. Immunohistochemical staining

Anesthetized mice (140 mg/kg of ketamine and 12 mg/kg of xylazine) were transcardially perfused with ice cold PBS followed by 4% paraformaldehyde (Electron Microscopy Science). Whole brains were dissected and postfixed overnight (4°C), followed by submersion in 30% sucrose for 48 hours. Coronal brain sections containing the hypothalamus were cut at 30 µm thickness with a freezing stage microtome (Leica SM2010R) and stored in freezing solution (phosphate buffer, 30% sucrose, 30% ethylene glycol) at −20 C until the day immunohistochemical (IHC) staining was performed. For the IHC, sections were washed with PBS, blocked with 2.5% normal donkey serum (Jackson ImmunoResearch Laboratories, catalog no. 017-000-121) and Triton X-100 0.5% for 1 h, and incubated overnight with rabbit anti-IBA1 (1:1000, FUJIFILM Wako), goat anti-IBA1 (1:1000, Novus), rat anti-CD68 (1:1000; Bio-Rad) and rabbit anti-POMC (1:4000; Phoenix Pharmaceuticals) polyclonal antibodies at 4°C. Sections were then incubated with the corresponding secondary antibodies, donkey anti-rabbit Alexa Fluor 488, donkey anti-goat Alexa Fluor 488, donkey anti-rat Alexa Fluor 555 and donkey anti-rabbit Alexa Fluor 647 (1:500, Thermo Fisher Scientific), for 1 h at room temperature. Stained sections were mounted on slides and cover slipped in the presence of mounting medium with DAPI for nuclear staining (Fluoromount-G; Thermo Fisher Scientific).

Images were captured on a Keyence BZ-X800 fluorescence microscope. Number of immunoreactive-positive IBA1 cells, POMC neurons, and the percent area occupied by IBA1 staining were determined per half section in a blinded fashion using Fiji (ImageJ Fiji; National Institutes of Health). Only anatomically matched section that included the mediobasal hypothalamus were used for quantification.

For microglial Sholl analysis, 60x z-stack images were captures on a Keyence BZ-X800 fluorescence microscope. In brief, three microglia per mouse were analysed in the arcuate nucleus (ARC) of the hypothalamus. Z-stack images were compiled into a 3D image, transformed into binary, and a fixed threshold was set to visualize microglia and processes. First, microglia cells were reconstructed using the Simple Neurite Tracer (SNT) plugin in Fiji. Then, Sholl analysis was performed using the Neuroanatomy Sholl plugin in Fiji. The start radius was set at 0.00 µm, step size of 1 µm and end radius was variable to cover all microglia processes. Number of intersections were automatically computed by the software.

To further characterize microglial morphology and quantify overlap between microglia and CD68 or POMC signals, confocal z-stack images were acquired using a Yokogawa W1 spinning disk confocal microscope (Nikon) equipped with a 100X plan apochromat silicon immersion objective at the Lynn & Mike Garvey Imaging Core, UW Institute for Stem Cell and Regenerative Medicine. Images were imported into Imaris software (Oxford Instruments) and the FilamentTracer module was applied to the IBA1 channel to determine filament length, number of branch points (total bifurcations), and branch endpoints (to determine arborization of microglial branches). Soma reconstruction was also performed to determine soma volume, ellipticity (oblate), and sphericity. Subsequently, intensity-base three-dimensional (3D) renderings were performed using the Surface module for the IBA1, CD68 and POMC channels. The volume of CD68 or POMC signals within 0 μm distance of the IBA1 signal was calculated to quantify overlap with microglia. All imaging and analyses for a given brain region were performed blinded and in randomized order, using identical acquisition and analysis settings.

### 2.12. Statistical analysis

Group sizes were determined using power calculations. Differences in the number of mice per group (genotype) resulted from the use of available littermate Ctls and KOs from our colony, and the exclusion of mice meeting pre-established criteria, including malocclusion or dermatitis. Littermate mice were randomized into experimental groups based on similar body weights. All analyses were performed in a blinded manner. To minimize potential confounding effects, measurements were performed in an alternating sequence of control and KO mice. Data are presented as mean ± SEM. Statistical analysis were conducted using GraphPad Prism. Normality was assessed for each dataset using the D’Agostino-Pearson omnibus test. For datasets that passed normality, unpaired two-tailed Student’s t-tests and two-way ANOVA followed by Sidak post hoc test were performed. For non-normally distributed data, Mann-Whitney and Dunn post hoc test were applied. Statistical outliers were identified by the ROUT (Q=1%) method. Probability (P)values of <0.05 were considered statistically significant.

## 3. Results

### 3.1. Validation of MG-ER_α_KO mouse model

To determine the impact of microglial estrogen signaling on metabolism, we generated a microglia-specific ERα knockout mouse model (Cx3cr1^CreER^::ERα^fl/fl^) using Cx3cr1^CreER^ to drive tamoxifen-inducible recombination of the floxed ERα alleles.

Hypothalamic genomic DNA amplification by PCR produced the expected _∼_250-bp Cre-mediated deletion fragment in MG-ERαKO mice but not in their littermate Ctl mice, confirming excision of the loxP-flanked ERα (*Esr1*) gene (Figure 1B). To determine the efficiency of knockdown specifically in microglia, gene expression analysis was performed in microglial primary cultures and in microglia isolated from adult brains of Ctl and MG-ERαKO mice. Tamoxifen-induced recombination in primary microglia cultures resulted in a 71% reduction of *Esr1* expression (Figure 1C). Similarly, microglia isolated from adult brains of MG-ERαKO exhibited ∼90% reduction in *Esr1* expression compared to Ctl mice (Figure 1D).

### 3.2. Microglial ER**_α_** deletion increases susceptibility to DIO in male but not female mice

To assess the effect of microglia-specific ERα deficiency in DIO, we monitored body weight, food intake, and fat mass in female and male MG-ERαKO and Ctl mice fed with HFD for 15 weeks (Figure 2A). While female MG-ERαKO and Ctl mice were indistinguishable in terms of body weight gain, food intake, and fat pad mass (Figure 2B-F), male MG-ERαKO showed accelerated weight gain during HFD feeding compared to Ctl mice (Figure 2G). Remarkably, the divergence in weight gain occurred within 3 weeks of HFD initiation (Figure 2G). Importantly, no difference in body weight was observed on chow diet in either sex (Supplemental Figure 1 A, C). A trend to lower cumulative food intake, measured in a group-housed setting, and increased fat mass was observed in MG-ERαKO male mice (Figure 2H-I); in contrast, there were no differences in lean mass (Figure 2J). Together these data support a role of microglia ERα signaling in protecting from DIO in male, but no female mice, by a mechanism independent of hyperphagia.

**Figure 2.**
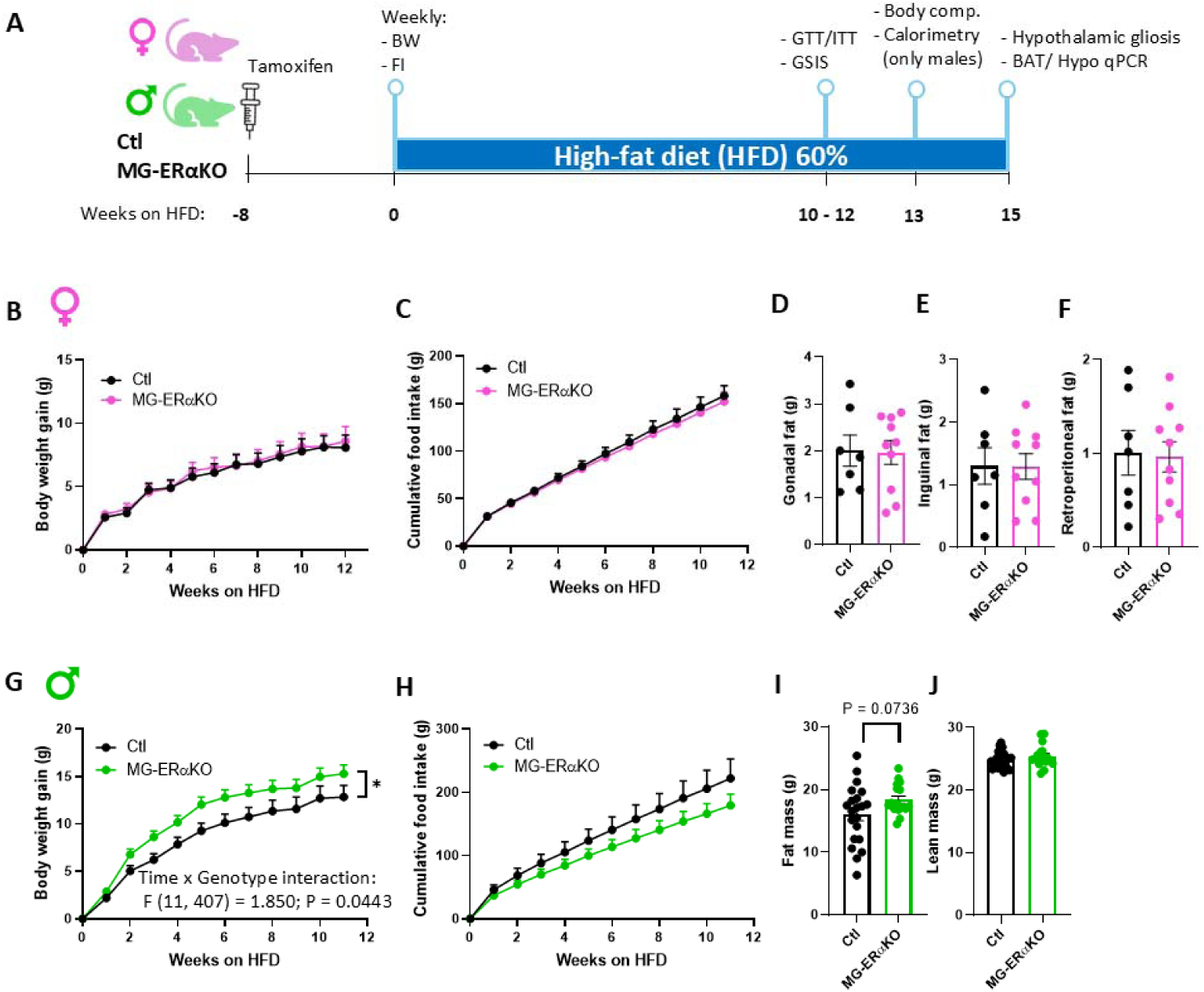
**Microglial ER**_α_ **deficiency exacerbates diet-induced obesity in a sex-dependent manner.** (A) Schematic illustration of the experimental design. (B-C and G-H) Body weight gain and cumulative food intake in MG-ERαKO female (B-C) and male (G-H) mice vs Control (Ctl) littermates recorded for 12 weeks of HFD feeding. (D-F) Gonadal (D), inguinal (E) and retroperitoneal (F) fat pads weight in female mice measured at study termination (15 weeks of HFD). (I-J) Total fat mass (I) and lean mass (J) in male mice measure by quantitative magnetic resonance spectroscopy at week 13 of HFD. Data are presented as mean ± SEM. Females Ctl n=19 and MG-ERαKO n=20; Males Ctl n=21 and MG-ERαKO n=18. A subset of females was used for fat pad measurements Ctl n=7 and MG-ERαKO n=10. Body weight and food intake data was analysed by Two-way ANOVA. Fat mass was analysed by Student’s *t*-test.

### 3.3. Male mice lacking microglial ER**_α_** have reduced energy expenditure during HFD feeding

To determine whether the excess weight gain observed in male MG-ERαKO mice was due to changes in energy expenditure, we performed indirect calorimety on Ctl and KO males after 13wk of HFD feeding. MG-ERαKO male mice exhibited a significant decrease in energy expenditure (EE) in both the dark and light cycles (Figure 3A-B) with no changes in respiratory quotient (RQ) (Figure 3C-D) or locomotor activity (Figure 3E-F). Consistent with the group housed data (Figure 2H), food consumption in a 24h-period, measured in metabolic cages (single-housed setting), was reduced in MG-ERαKO males relative to Ctls (Figure 3G-H). Food intake tended to be lower both in the dark and light cycles, although the differences did not reach statistical significance (Figure 3H). Given the higher body weight of MG-ERαKO mice, the decreased food intake likely represents an inadequate compensatory response to the lower energy expenditure induced by the loss of microglia ERα signaling. To support this hypothesis, gene expression analysis performed in brown adipose tissue (BAT) revealed decreased expression of thermogenic genes cell death-inducing DFFA-like effector A (*Cidea*) and carnitine palmitoyltransferase 1B (*Cpt1b*) in KO mice (Figure 3I), consistent with decreased BAT thermogenesis. Interestingly, stearoyl-CoA Desaturase1 (*Scd1*) expression was significantly elevated in MG-ERαKO male mice (Figure 3I), which is associated with increased body weight gain and fat mass [52].

**Figure 3.**
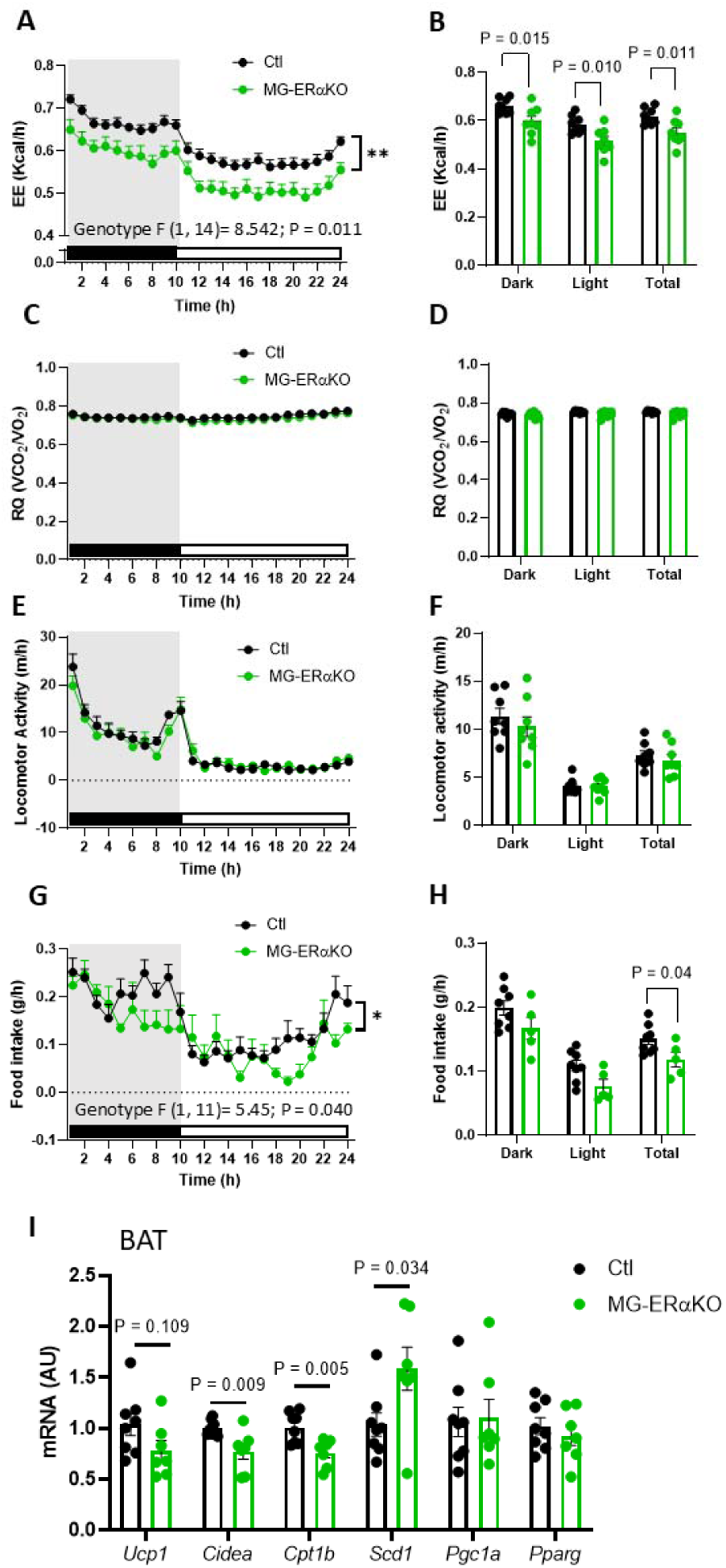
**Loss of microglial ER**_α_ **decreases energy expenditure in male mice.** Energy expenditure (EE; A), respiratory quotient (RQ; C), locomotor activity (E) and food intake (G) represented as time-course profile in MG-ERαKO and Ctl male mice fed with a HFD for 13 weeks (A, C, E, G; grey area represents the dark cycle). Binned dark phase, light phase, and 24h total averages for EE (B), RQ (D), locomotor activity (F) and food intake (H). (I) Brown adipose tissue (BAT) mRNA expression normalized to values from Ctl mice. Data from metabolic cages are presented as mean ± SEM of 4 consecutive days of measurements. Males Ctl n=8 and MG-ERαKO n=8, except for food intake where MG-ERαKO n=5. Time course data was analysed by two-way ANOVA, while averages of dark, light, and 24h total by student’s *t*-test. BAT gene expression data are presented as mean ± SEM of 7-8 mice per group and analysed by student’s *t*-test.

### 3.4. Loss of microglial ER**_α_** signaling during HFD impairs insulin sensitivity

To evaluate the effects of microglia ERα deficiency on glucose homeostasis, we first performed an i.p. GTT in chow-fed mice and found no differences between MG-ERαKO and Ctl mice in either sex (Supplemental Figure 1 B, D). Under DIO, we then assessed glucose regulation by conducting i.p. GTT, ITT, and glucose-stimulated insulin secretion (GSIS) tests on MG-ERαKO and Ctl mice of both sexes. Equivalent glucose tolerance was observed in both genotypes and both sexes. In contrast, both male and female MG-ERαKO mice exhibited mild insulin resistance (Figure 4A-B and F-G). GSIS test resulted in no differences of plasma insulin levels at 0 and 15 min after glucose administration between genotypes in females (Figure 4C). While blood glucose at 15 min was significantly elevated in MG-ERαKO females during the GSIS test, HOMA-IR remained unchanged (Figure 4D-E). In males, there was a significant increase in insulin levels 15 min after glucose injection in MG-ERαKO compared to Ctl mice (Figure 4H). Despite the elevated insulin response, blood glucose levels remained comparable between genotypes at 15 min (Figure 4I), consistent with insulin resistance. This conclusion was supported by the significant increase in HOMA-IR (Figure 4J). Together, these findings suggest that ERα signaling in microglia is required to limit obesity-associated insulin resistance in both sexes. While in males, the insulin resistance of MG-ERαKO mice may result from the increased fat mass, the underlying weight-independent mechanism in females remains unclear.

**Figure 4.**
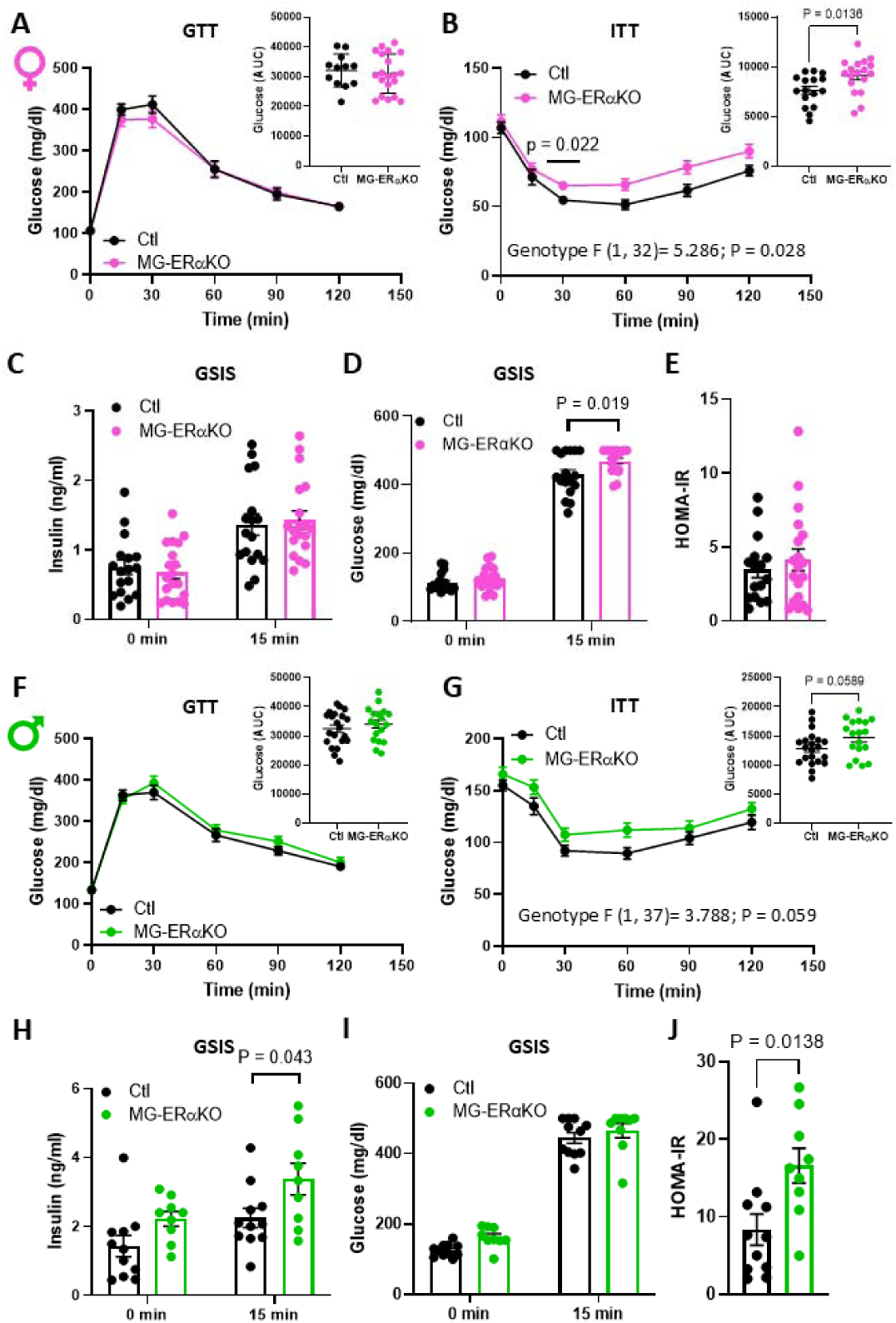
**Deletion of microglial ER**_α_ **reduces insulin sensitivity in males and females.** (A, F) Glucose tolerance test (GTT; 2 g/kg ip), (B, G) insulin tolerance test (ITT; 0.75 U/kg ip), and (C-D, H-I) glucose-stimulated insulin secretion (GSIS, 2 g/kg ip) represented as insulin and glucose levels at time = 0 min and time = 15 min after glucose administration in MG-ERαKO female (A-D) and male (F-I) mice vs controls (Ctl). Insets in A, B, F, G represent area under the curve (AUC) glucose. (E, J) Homeostatic model assessment of insulin resistance (HOMA-IR) index in female (E) and male (J) mice. Data are presented as mean ± SEM. GTT, ITT and GSIS in females included Ctl n=19 and MG-ERαKO n=20; GTT and ITT in males included Ctl n=21 and MG-ERαKO n=18 and GSIS included Ctl n=11 and MG-ERαKO n=9. Two-way ANOVA followed by Sidak’s post hoc test was used for GTT and ITT analysis, and Student’s *t*-test for AUC comparisons.

### 3.5. Microglial ER**_α_** deficiency promotes microglial activation during HFD feeding

Male rodents consuming a HFD display a hypothalamic injury response characterized by the accumulation of activated pro-inflammatory microglia [22,23]. In contrast, females are largely resistant to these changes [31], raising the possibility that microglia ERα signaling limits HFD-induced microglial activation. Therefore, we assessed microglial number, immunostaining density, and morphometry in the arcuate (ARC) nucleus of the hypothalamus in male and female MG-ERαKO and Ctl mice using the microglial marker IBA1. In males, there were no differences in IBA1^+^ cell number or percentage area in the ARC between genotypes (Figure 5A-C), but the number of intersections and the AUC calculated from Sholl analysis were both significantly reduced (Figure 5D-F). These data suggest that male microglia lacking ER signaling assumed a more activated profile, characterized by shortened and retracted branches. Consistent with these findings, filament length was also decreased though no changes were found in the total number of ramifications (branch points and endpoints) (Figure 5G-J). Microglial soma volume and sphericity were unchanged between genotypes (Figure 5K-L), whereas ellipticity was significantly reduced in KO mice (Figure 5M), suggesting an intermediate or atypical activation profile rather than a robust, pro-inflammatory response. In contrast, microglial ERα deficiency in female mice led to an increased number of IBA1^+^ cells and elevated IBA1 expression in the ARC (Supplemental Figure 2A-C), approaching levels typically observed in males (Figure 5A-C). However, no morphological differences were detected between genotypes in the microglial Sholl analysis (Supplemental Figure 2D-F) nor were there changes in filament length, branching complexity, soma volume or sphericity (Supplemental Figure 2H-L). Nevertheless, soma ellipticity was significantly reduced in MG-ERαKO (Supplemental Figure 2M), mirroring the effect seen in male KOs (Figure 5M). Despite these alterations in microglial morphology, no differences were detected in inflammatory gene expression in either whole hypothalamic tissue or isolated hypothalamic microglia from both sexes (Supplemental Figure 3A-B and 3E-F). Collectively, these findings support the hypothesis that loss of microglial ERα signaling predisposes male mice to HFD-induced microglial activation in the ARC.

**Figure 5.**
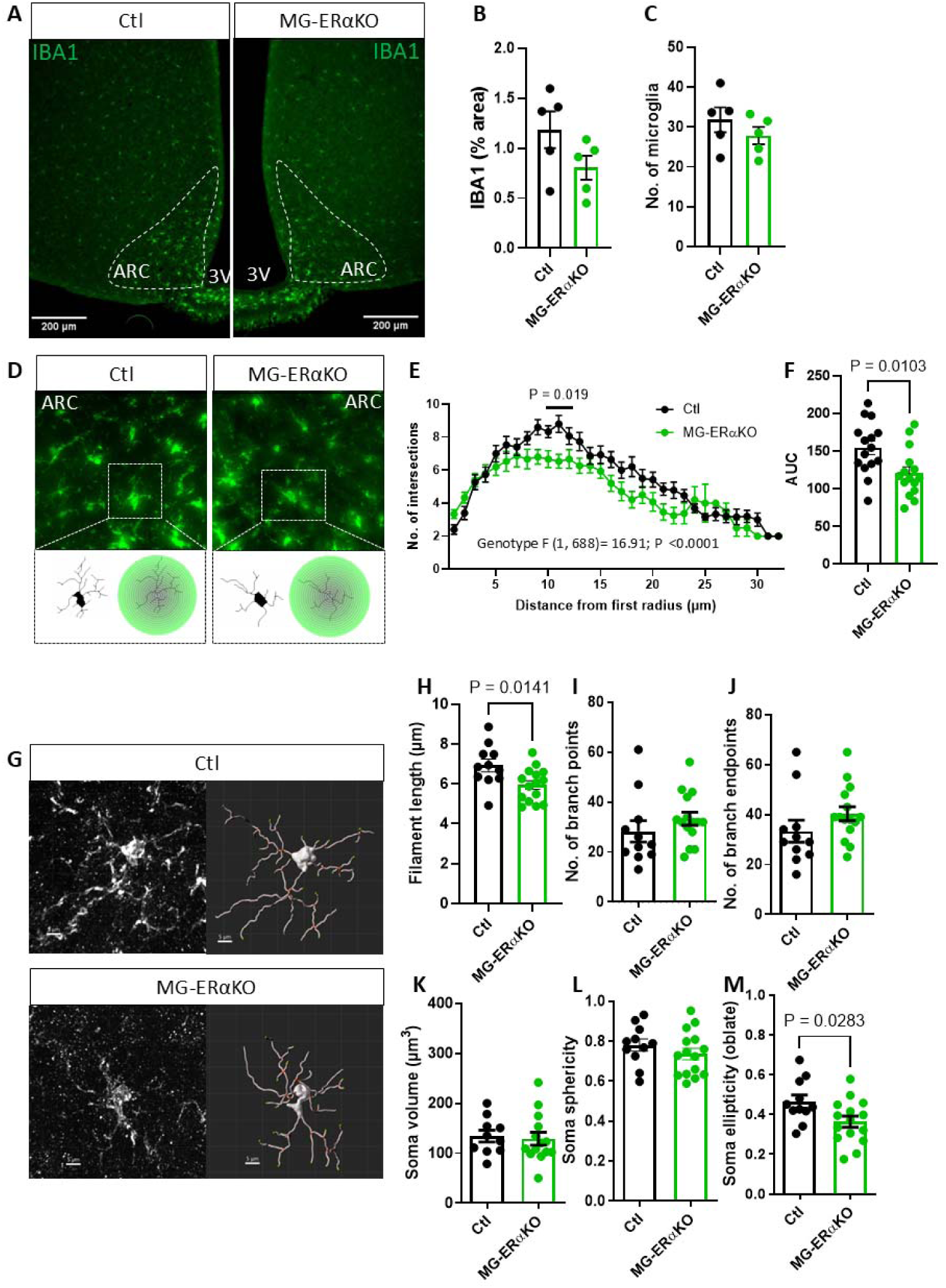
**Microglial ER**_α_ **deletion promotes hypothalamic microglial activation in HFD-fed male mice.** (A) Representative images of IBA1 immunostaining in the arcuate nucleus (ARC) of the hypothalamus from MG-ERαKO male mice and controls (Ctl) after 15 weeks of HFD. 3v = third ventricle (B-C) Quantification of (B) IBA1 % area and (C) the number of microglia (IBA1-positive cells) per half-section. (D) Representative images of high magnification IBA1-labeled microglia in the ARC from Ctl and MG-ERαKO male mice, with insets showing reconstructed microglia and concentric radii from the soma center. (E) Sholl analysis quantification, represented as the number of microglial process intersections per radius, and (F) the corresponding AUC. (G) Confocal images and 3D reconstruction of ARC microglia from Ctl and MG-ERαKO male mice. (H-M) Morphometric analysis of reconstructed microglial filaments and soma: (H) filament length, (I) number of branch points (bifurcations), (J) number of branch endpoints (microglial complexity), (K) soma volume, (L) soma sphericity, and (M) soma ellipticity. Data are presented as mean ± SEM. Sholl and morphological analysis were performed as follows (Ctl: n = 4 mice, 12 microglia cells; MG-ERαKO n = 5 mice, 15 microglia) in ARC sections. Statistical significance was determined by Student’s t-test vs Ctl group. Two-way ANOVA followed by Sidak’s post hoc test was used for Sholl analysis, and Student’s *t*-test was applied for AUC comparison.

### 3.6. Microglia ER**_α_** KO male mice have reduced POMC neurons in the arcuate nucleus

Current evidence suggests that DIO is associated with dysfunction of POMC neurons, a hypothalamic cell population that maintains energy homeostasis by modulating food intake and energy expenditure [17–19]. We and others have demonstrated that changes in microglial signaling alter POMC neuron properties in HFD-fed mice, including electrophysiological activity, neuron-microglia interactions, and downstream melanocortin signaling [20,24–26]. In this study, we quantified the percentage of microglial cell volume occupied by CD68 (a marker of phagocytosis), the percentage of microglial volume overlapping POMC neurons, and the number of POMC neurons in the ARC. The percentage of overlapping volume between CD68 and IBA1 tended to be higher in MG-ERαKO male mice compared to controls (p=0.07) (Figure 6A-B). The percentage volume of microglia overlapping with POMC neurons was significantly greater in MG-ERαKO male mice than in Ctl (Figure 6C-D). Despite no differences in hypothalamic mRNA levels of neuropeptides (AgRP, NPY, POMC and CART) in either sex as assessed by qPCR (Supplemental Figure 3C-D), MG-ERαKO male mice exhibited a reduced number and volume of POMC neurons in the MBH as determined by IHC (Figure 6E-G). These data suggest that the reduction of POMC neurons in MG-ERαKO mice may result from increased microglial phagocytic activity.

**Figure 6.**
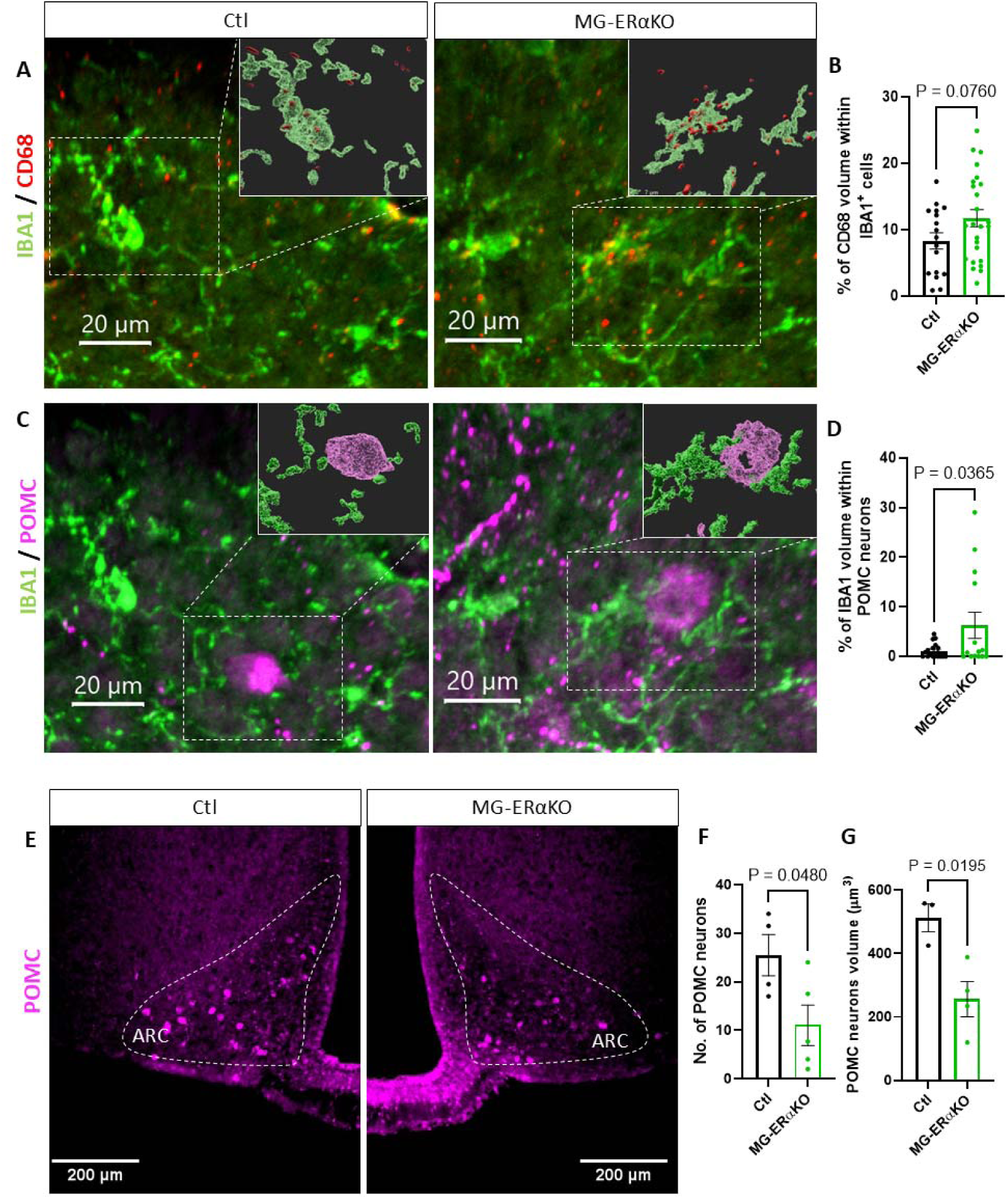
**Microglial-POMC interactions are increased and POMC neurons number reduced in HFD-fed male mice lacking ER**_α_. (A) Representative confocal images of microglial IBA1 and CD68 immunostaining, with insets showing 3D reconstructions of IBA1-CD68 colocalization in the ARC from Ctl and MG-ERαKO male mice. (B) Quantification of the percentage of microglial (IBA1^+^ cell) volume occupied by CD68 staining (measure of phagocytic activity). (C) Representative images of IBA1 and POMC immunostaining with insets displaying 3D reconstructions of IBA1-POMC colocalization in the ARC from Ctl and MG-ERαKO male mice. (D) Quantification of the percentage of POMC neuron volume occupied by IBA1 staining (measure of microglial-POMC neuron contact). (E) Representative images of POMC immunostaining in the ARC and quantification shown as (F) number of POMC neurons per half-section and (G) POMC volume in MG-ERαKO and Ctl male mice. Data are presented as mean ± SEM. Colocalization analysis performed as follows (n = 3 mice, 18 microglia in Ctl; n = 4 mice, 24 microglia cells in MG-ERαKO). Statistical significance was determined by Student’s t-test.

## 4. Discussion

The female sex steroid E2 has pleiotropic reproductive and metabolic effects in both females and males. Through its primary receptor ERα, E2 is known to reduce obesity susceptibility via actions in the CNS including several neuronal and non-neuronal populations of the hypothalamus [7,12,53,54]. However, little is known about the role of estrogen signaling in microglia, a cell type that has been implicated in the pathogenesis of obesity [20,22–25,30,31]. This study provides the first evidence that microglia-specific ablation of ERα promotes morphologic alterations to hypothalamic microglia toward more activated profiles while exacerbating obesity-associated insulin resistance during HFD feeding in both males and females. In males, loss of microglial estrogen signaling is also associated with a reduced number of hypothalamic POMC neurons and increased HFD-induced weight gain as a result of lower energy expenditure. Together, these studies reveal a sex-specific cell autonomous role of ERα signaling to limit microglial activation, weight gain, and insulin resistance during HFD feeding.

Several studies have shown an inverse correlation between metabolic disease and reproductive capacity [1,48], highlighting the beneficial effects of estrogen signaling in females [1,55]. Moreover, young female mice are resistant to DIO and HFD-induced MBH microglial activation [31,56–58]. However, estrogen deficiency (e.g. by OVX) eliminates these protections, leading to male level gliosis and DIO, effects reversed by estrogen replacement therapy [37,38]. These findings support the hypothesis that a direct effect of estrogen on microglia limits HFD-induced gliosis and obesity. To address this possibility, we analysed the metabolic and histopathologic phenotypes of an inducible microglia-specific ERα knockout mouse fed with HFD. Contrary to our hypothesis, we observed that female MG-ERαKO mice demonstrated only mild insulin resistance and minor cell morphology changes in microglia without increased weight gain, suggesting ERα-mediated protection against DIO in females largely involves other CNS cell types. This is consistent with prior studies showing that the anorexigenic and hypermetabolic effects of E2 in females, but not males, are mainly regulated by ERα-positive POMC neurons in the ARC and ERα-positive SF1 neurons in the ventromedial hypothalamus (VMH), respectively [12,59]. More recently, studies have also demonstrated that E2 potentiates the anorectic actions of leptin by Cited1-expressing POMC neurons [53], while Mc4r neurons in the VMH mediates the E2 effect on thermogenesis and physical activity [60]. Thus, increased MBH microgliosis observed in OVX females fed with HFD is likely a consequence of altered neuronal activity and not directly causative of DIO susceptibility.

Although estrogens are classically viewed as more relevant to the biology of females with levels much higher than in males, ERα genetics and induced mutations in mouse models have revealed critical biological and pathological roles for estrogen in both sexes [61]. In the context of metabolic diseases, ERα mutations in men and in male mice predispose to obesity [13,62]. However, pan-neuronal deletion of ERα induces only mild weight gain in males while specific ERα deletion in POMC or SF1 neurons has no weight impact [12]. Here, we show that ERα deletion in microglia is sufficient to exacerbate DIO in male mice through a marked reduction of energy expenditure. This result contrasts with most previous studies involving microglial contributions to metabolism in which genetic modifications (e.g. ablation of the NF-κB pathway, prostaglandin signaling, mitochondrial uncoupling proteins, clock genes, etc.) have resulted in altered food intake rather than energy expenditure [24,25,30,35]. Interestingly, analysis of BAT revealed a downregulation of thermogenic genes (*Cidea* and *Cpt1*) and an upregulation of *Scd1*, a monounsaturated fatty acid synthesis gene that promotes obesity [52]. Recently, *Scd1* has also been linked with sympathetic activity and BAT thermogenesis, suggesting a potential role in altering energy expenditure [63].

Insulin resistance and glucose intolerance typically track with the degree of adiposity. Nevertheless, MG-ERαKO female mice develop mild insulin resistance despite normal body weight, suggesting that microglial estrogen signaling directly modulates glucose homeostasis. This idea is supported by a recent study demonstrating microglial inflammation paradoxically improves glucose tolerance while promoting weight gain [50]. Notably, male MG-ERαKO mice also display mild insulin resistance, but this likely results from their increased adiposity.

Microglial reactivity during acute and chronic interventions is sexually dimorphic in certain contexts [64]. For example, during HFD feeding of male mice, MBH microglial number increases with cells assuming a more activated morphology; in contrast, females do not show this response [31]. Similarly, we find that ablating microglial ERα promotes a transition toward less ramified, more ameboid microglial morphology in the MBH of male mice. Consistent with this observation, microglial ERα deficiency tends to increase microglial CD68 levels (a marker of phagocytosis) and contact between microglia and POMC neurons. Ultimately, these alterations to microglial properties are associated with a reduced number of POMC neurons in the arcuate nucleus, a possible mechanism to explain the increased adiposity in the KO mice. This is consistent with previous reports showing that HFD feeding induces alterations in MBH POMC neurons, including synaptic remodeling, reduced neuronal activity, and neuronal loss [18,20,22,26]. Overall, these findings support a model in which maintenance of microglial estrogen signaling in males limits HFD-induced disruption of POMC neuron integrity during HFD feeding. This mechanism is consistent with our previous report of DIO protection in a mouse lacking the PGE2 receptor EP4 in microglia, a phenotype associated with reduced microglial CD68 expression and phagocytic capacity along with preserved POMC neuron architecture [25].

In summary, this study provides further clarity on estrogen action in specific cell populations of the CNS that contribute to the metabolic differences between males and females. Specifically, we uncovered a previously unrecognized role for microglial ERα signaling in limiting diet-induced obesity in males. While estrogen’s protective metabolic effects in females primarily involve hypothalamic neurons, our findings demonstrate that in males, microglial ERα restrains diet-induced gliosis, preserves POMC neuron integrity, and sustains energy expenditure through brown adipose tissue thermogenesis. These results highlight microglia as critical mediators of CNS estrogen action and point to glial pathways as potential therapeutic targets for metabolic disease.

## CRediT authorship contribution statement

**Inmaculada Velasco**: Conceptualization, Data curation, Formal analysis, Investigation, Methodology, Writing – original draft. **Jeremy M. Frey**: Investigation, Methodology, Validation. **Vladislav Baglaev**: Formal analysis, Investigation. **Tristan Jafari**: Investigation. **Thomas Huang**: Investigation. **Sophia Schwie**: Investigation. **Olivia Santiago**: Investigation, Methodology. **Rachael D. Fasnacht**: Investigation, Methodology. **Joshua P. Thaler**: Conceptualization, Funding acquisition, Resources, Writing – review and editing. **Mauricio D. Dorfman**: Conceptualization, Funding acquisition, Investigation, Methodology, Project administration, Resources, Supervision, Writing – review and editing.

**Declaration of generative AI and AI-assisted technologies in the writing process** During the preparation of this manuscript, the authors used ChatGPT (OpenAI, 2025) to improve the clarity and readability of the text. Following the use of these tools, the authors carefully reviewed and edited the content and take full responsibility for all aspects of the manuscript. Graphical abstract made with a licensed version of BioRender.com.

---

**Supplemental Figure 1.**
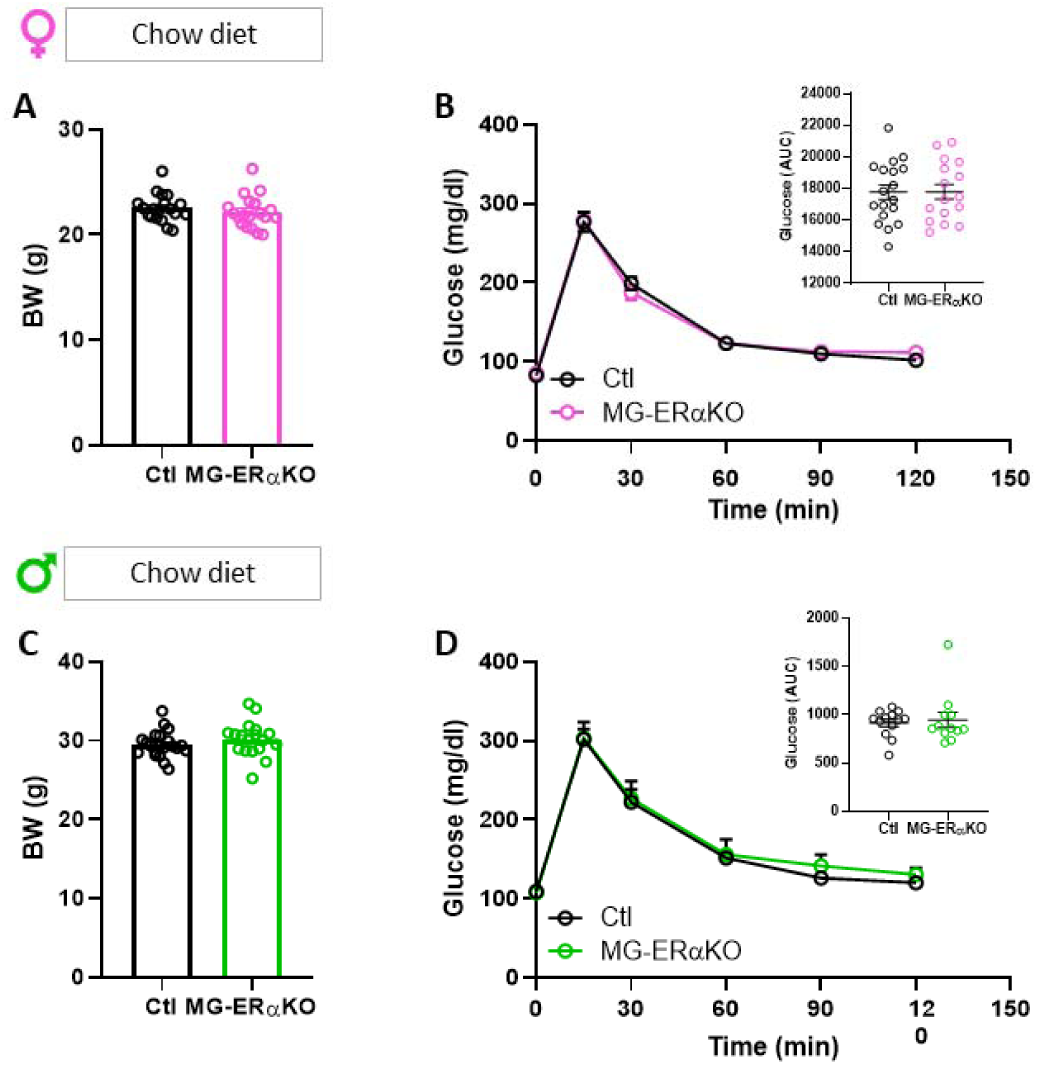
E**R**α **signaling in microglia does not impact body weight gain and glucose tolerance in chow-fed mice.** (A) Body weight in female and (C) male mice fed with chow diet. Females Ctl n = 19 and MG-ERαKO n=20; Males Ctl n = 21 and MG-ERαKO n=18. (D) Intraperitoneal glucose tolerance test (GTT; 2 g/kg ip) in female and (E) male mice under chow diet feeding. Insets in D and E represent the area under the curve (AUC) for blood glucose. Females Ctl n = 18 and MG-ERαKO n = 16; Males Ctl n = 12 and MG-ERαKO n = 12. Data are presented as mean ± SEM.

**Supplemental Figure 2.**
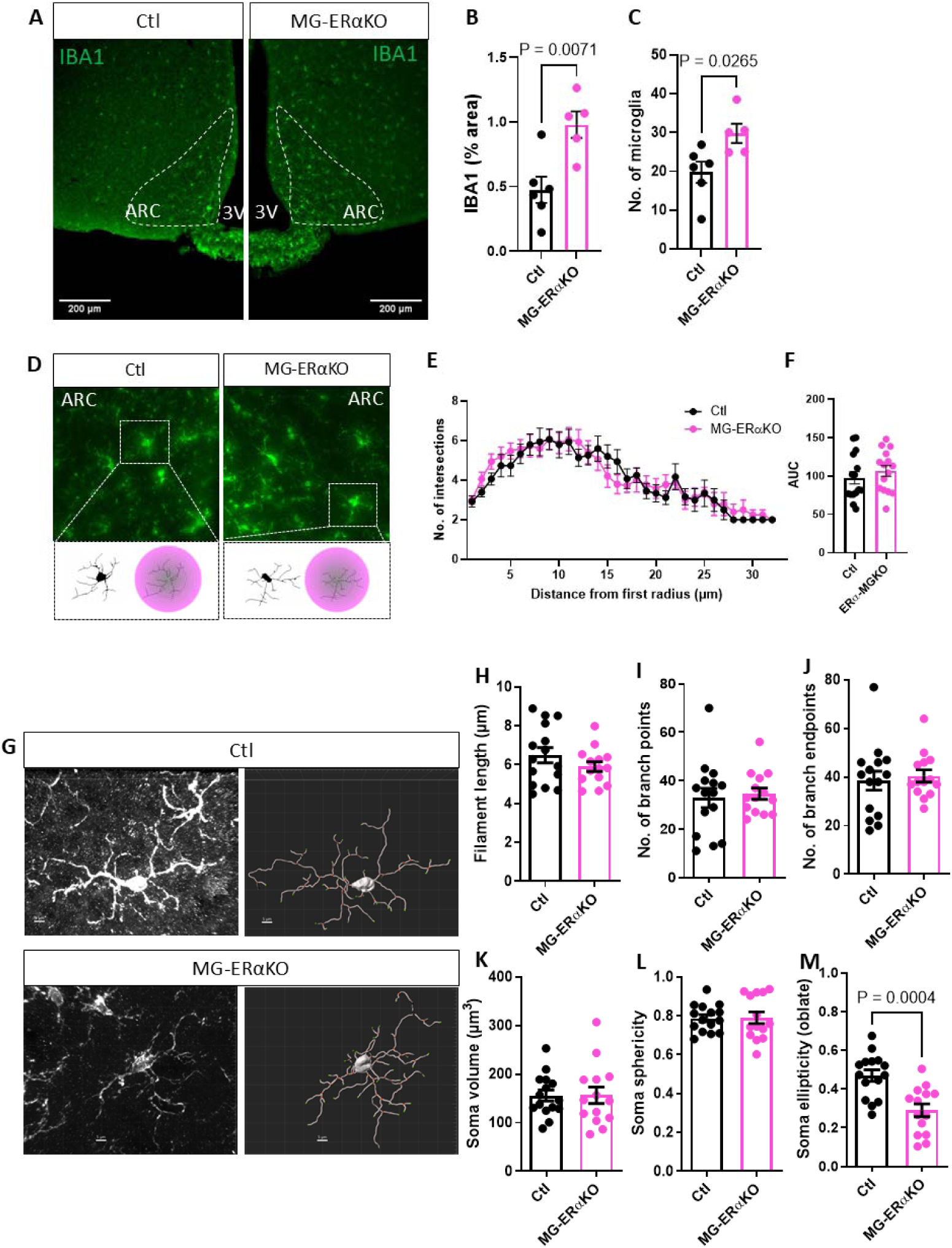
Microglial ER_α_ deletion does not alter microglial morphology in female mice fed a HFD. (A) Representative images of IBA1 immunostaining in the arcuate nucleus (ARC) of the hypothalamus from MG-ERαKO female mice and controls (Ctl) after 15 weeks of HFD. 3v = third ventricle (B-C) Quantification of (B) IBA1 % area and (C) the number of microglia (IBA1-positive cells) per half-section(E) Sholl analysis quantification, represented as the number of microglial process intersections per radius, and (F) the corresponding AUC. (G) Confocal images and 3D reconstruction of ARC microglia from Ctl and MG-ERαKO male mice. (H-M) Morphometric analysis of reconstructed microglial filaments and soma: (H) filament length, (I) number of branch points (bifurcations), (J) number of branch endpoints (microglial complexity), (K) soma volume, (L) soma sphericity, and (M) soma ellipticity. Data are presented as mean ± SEM. Sholl and morphological analysis were performed as follows (n = 5 mice, 15 microglia in Ctl; n = 5 mice, 15 microglia in MG-ERαKO) in ARC sections. Statistical significance was determined by Student’s t-test.

**Supplemental Figure 3.**
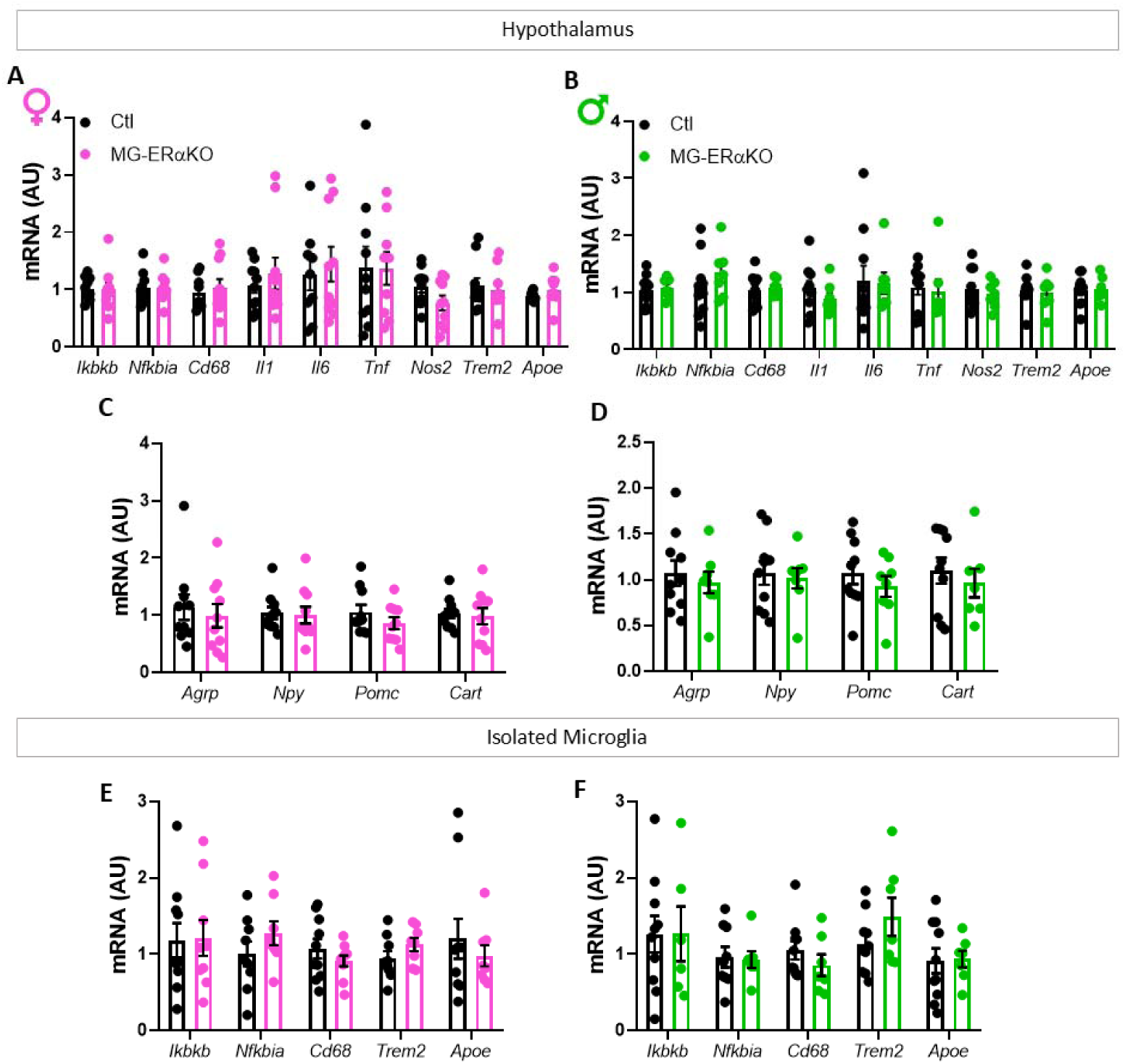
Whole hypothalamic and isolated hypothalamic microglial gene expression. (A) Expression of hypothalamic neuropeptides involved in food intake and energy homeostasis, and (B-C) expression of inflammatory genes in (B) whole hypothalamus and (C) isolated hypothalamic microglia from MG-ERαKO and control (Ctl) female (magenta) and male (green) mice. Data are presented as mean ± SEM.

