## Supplemental Figures for "Ablation of microglial estrogen receptor alpha predisposes to diet-induced obesity in male mice"

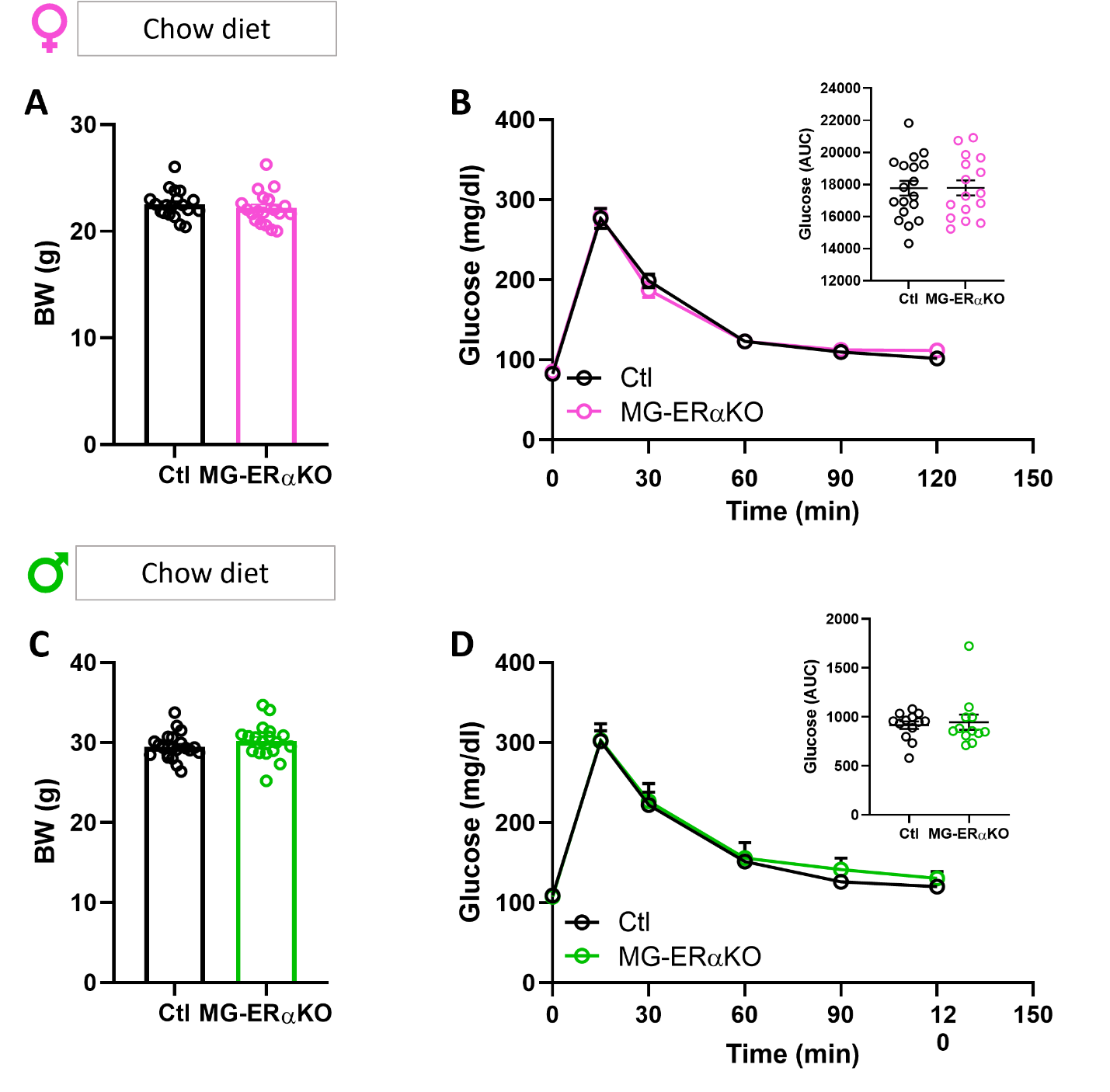

**Supplemental Figure 1. ERα signaling in microglia do not impact body weight gain and glucose tolerance in chow-fed mice.** (A) Body weight in female and (C) male mice fed with chow diet. Females Ctl n = 19 and MG-ERαKO n=20; Males Ctl n = 21 and MG-ERαKO n=18. (D) Intraperitoneal glucose tolerance test (GTT; 2 g/kg ip) in female and (E) male mice under chow diet feeding. Insets in D and E represent the area under the curve (AUC) for blood glucose. Females Ctl n = 18 and MG-ERαKO n = 16; Males Ctl n = 12 and MG-ERαKO n = 12. Data are presented as mean ± SEM.

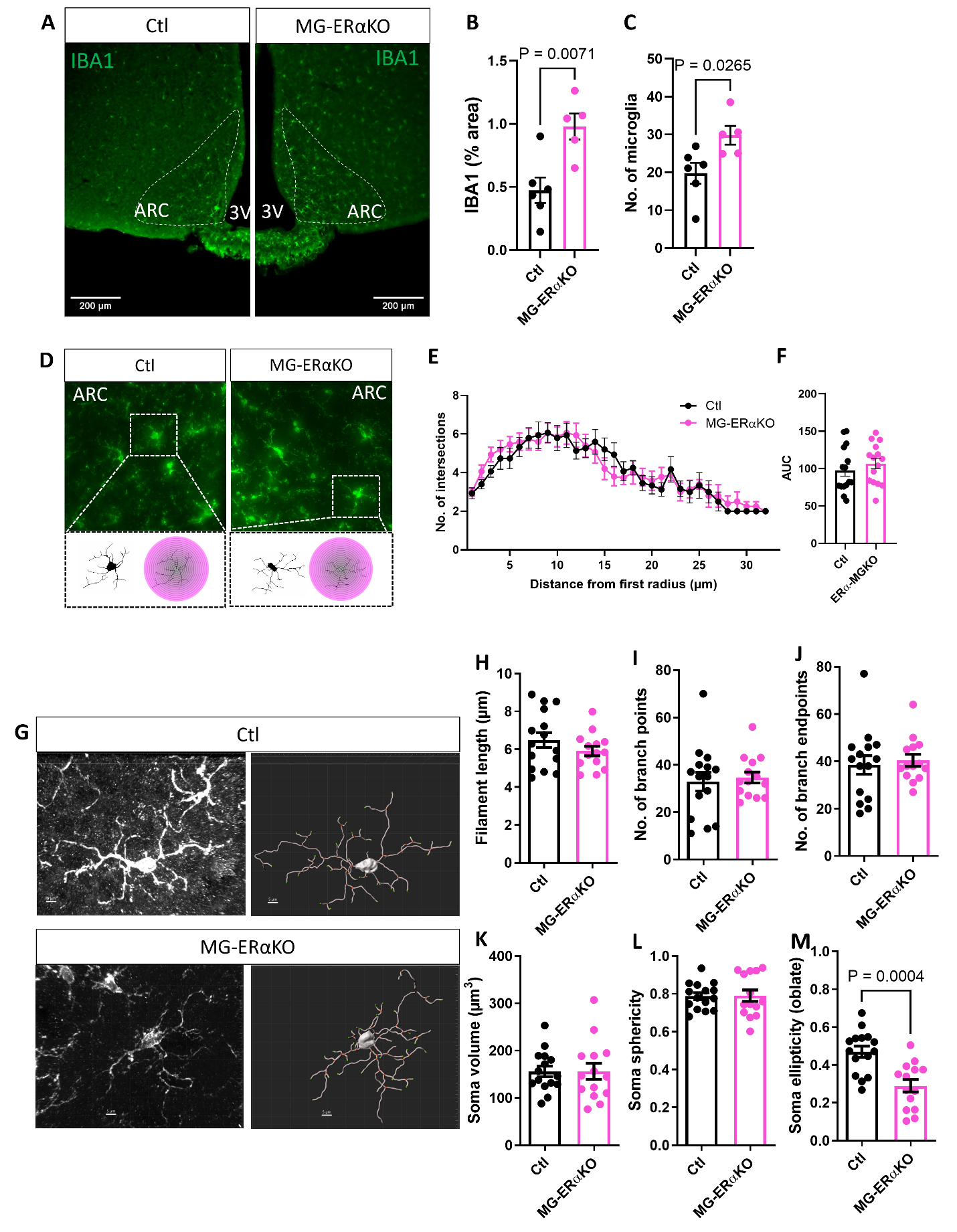

**Supplemental Figure 2. Microglial ERα deletion does not alter microglial morphology in female mice fed a HFD.** (A) Representative images of IBA1 immunostaining in the arcuate nucleus (ARC) of the hypothalamus from MG-ERαKO female mice and controls (Ctl) after 15 weeks of HFD. 3v = third ventricle (B-C) Quantification of (B) IBA1 % area and (C) the number of microglia (IBA1-positive cells) per half-section(E) Sholl analysis quantification, represented as the number of microglial process intersections per radius, and (F) the corresponding AUC. (G) Confocal images and 3D reconstruction of ARC microglia from Ctl and MG-ERαKO male mice. (H-M) Morphometric analysis of reconstructed microglial filaments and soma: (H) filament length, (I) number of branch points (bifurcations), (J) number of branch endpoints (microglial complexity), (K) soma volume, (L) soma sphericity, and (M) soma ellipticity. Data are presented as mean ± SEM. Sholl and morphological analysis were performed as follows (n = 5 mice, 15 microglia in Ctl; n = 5 mice, 15 microglia in MG-ERαKO) in ARC sections. Statistical significance was determined by Student’s t-test.

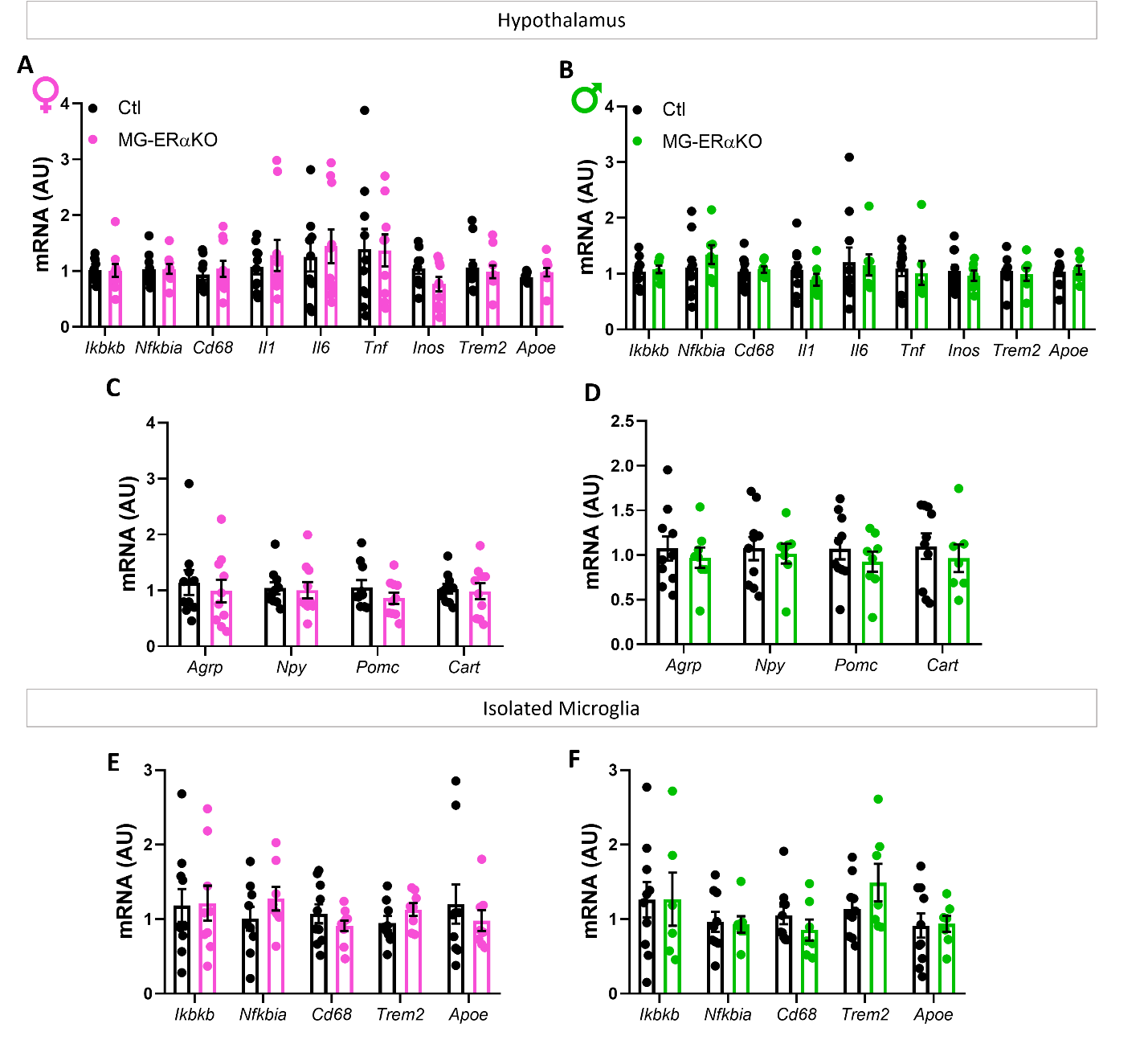

**Supplemental Figure 3. Whole hypothalamic and isolated hypothalamic microglial gene expression.** (A) Expression of hypothalamic neuropeptides involved in food intake and energy homeostasis, and (B-C) expression of inflammatory genes in (B) whole hypothalamus and (C) isolated hypothalamic microglia from MG-ERαKO and control (Ctl) female (magenta) and male (green) mice. Data are presented as mean ± SEM.
